# cFIT: Integration and transfer learning of single cell transcriptomes, illustrated by fetal brain cell development

**DOI:** 10.1101/2020.08.31.276345

**Authors:** Minshi Peng, Yue Li, Brie Wamsley, Yuting Wei, Kathryn Roeder

## Abstract

Large, comprehensive collections of scRNA-seq data sets have been generated that allow for the full transcriptional characterization of cell types across a wide variety of biological and clinical conditions. As new methods arise to measure distinct cellular modalities, a key analytical challenge is to integrate these data sets or transfer knowledge from one to the other to better understand cellular identity and functions. Here, we present a simple yet surprisingly effective method named *cFIT* for capturing various batch effects across experiments, technologies, subjects, and even species. The proposed method models the shared information between various data sets by a common factor space, while allowing for unique distortions and shifts in gene-wise expression in each batch. The model parameters are learned under an iterative non-negative matrix factorization (NMF) framework and then used for synchronized integration from across-domain assays. In addition, the model enables transferring via low-rank matrix from more informative data to allow for precise identification in data of lower quality. Compared to existing approaches, our method imposes weaker assumptions on the cell composition of each individual data set, however, is shown to be more reliable in preserving biological variations. We apply cFIT to multiple scRNA-seq data sets of developing brain from human and mouse, varying by technologies and developmental stages. The successful integration and transfer uncover the transcriptional resemblance across systems. The study helps establish a comprehensive landscape of brain cell type diversity and provides insights into brain development.

## 1 Introduction

Individual single-cell RNA sequencing (scRNA-seq) experiments have already been used to discover new cell states and reconstruct cellular differentiation trajectories. Through global efforts such as the Human Cell Atlas, researchers are now generating large, comprehensive collections of scRNA-seq data sets that profile a diverse range of cellular functions, which promise to enable high-resolution insight into processes underlying fundamental biology and disease. Multiple data sets and measurement assays of the same biological system are now widely available that reflect a similar set of biological processes. Specifically, experiment design with different technology and sample selection brought in exclusive coverage and strength of each data set. For instance, some experiment prepared samples covering a wide developmental stage while another focus on one particular stage. Besides, technologies such as Drop-seq, inDrop, and 10 × Genomics allow profiling with high throughput and low cost, enabling capturing rare populations despite limited sequencing depth, while other methods enable higher sensitivity, accuracy, and coverage. Therefore, assembling large, unified reference data sets from multiple sources can potentially bring unprecedented benefits.

Recent studies have shown that cellular features can be preserved across experimental systems from related biological contexts (Luecken and Theis, 2019). The information learned from different data sources can improve the analysis and interpretation of diverse biological systems. However, the advantages of integrated data can be compromised by differences due to experimental batch (experimental labs), sampling (sample acquisition and handling, sample composition, reagents or media, and sampling time) or technology (sequencing depth, sequencing lanes, read length, plates or flow cells, protocol) (Luecken et al., 2020). The challenge is exacerbated when technical differences in data sources are confounded with biological heterogeneity. Many methods have been established to integrate scRNA-seq studies across multiple experiments. Some methods employ supervised cross-domain transfer learning (Kiselev et al., 2018; Pliner et al., 2019; Ge et al., 2020) to remove domain effects with models learned from labeled data sets. These methods rely heavily on data labeling, thus failing to capture novel cell types and continuous trajectories. In contrast, unsupervised methods are less restrictive, and therefore more widely applied to integrate data from multiple resources (Butler et al., 2018; DePasquale et al., 2019; Hie et al., 2019; Stuart et al., 2019; Welch et al., 2019; Li et al., 2020). However, many of these methods tend to prioritize uniformity of mixing across different batches over preserving biological variation (Luecken et al., 2020). Such a principle can lead to a loss in biological heterogeneity and interpretability, especially when integrating collections of data sets with considerable differences in cellular composition. Besides, most existing methods work on the assumption that all data sets share most cell types, or that the within-domain biological variance defining distinct cell types dominates the cross-domain effects (Haghverdi et al., 2018; Stuart et al., 2019; Li et al., 2020); such assumptions do not hold when integrating biologically heterogeneous data sets or data consisting of continuously transitioning cell types or refined subtypes.

Here we present a simple but surprisingly effective unsupervised integration and transfer learning model, called cFIT, for Common Factor Integration & Transfer learning. The model assumes a shared common factor space across data sets, but with distortions and shifts on gene-wise expression unique by domain. Our model is motivated from the machine-learning subdomain of transfer learning (Raina et al., 2007; Pan and Yang, 2009; Vilalta and Drissi, 2002; Donahue et al., 2014; Tripuraneni et al., 2020), assuming that information is shared across different tasks, and common data representations can be learned and generalized to other unseen tasks. In this framework, the shared latent space represents the underlying biological processes across systems, such as common cell type compositions and developmental trajectories across measurements, samples, or even species. Once the robustness of a biological process is established, these learned latent spaces enable varied learning tasks across data platforms, modalities, and studies, through transfer learning.

The proposed model is capable of capturing various batch effects and integrating across various domains by employing a linear model that is far more parsimonious than existing methods such as LIGER (Welch et al., 2019). cFIT is powerful because it corrects for technical variation but does not remove biological heterogeneity, providing both flexibility and interpretability. For implementation, we derived an algorithm for inferring model parameters under an iterative non-negative matrix factorization (NMF) framework. The algorithm is also applicable for synchronized integration of across-domain assays. Besides, the learned biological signatures can be transfer learning to allow for precise inference for data with lower quality or smaller sample size.

Using cFIT, we successfully integrated two independent data sets derived from the developing human cortex (Nowakowski et al., 2017; Polioudakis et al., 2019). The integration disentangled the domain-specific technical effects with the biological processes unique to each data set, where the later is preserved and depicted in the recovered developing trajectories. The learned latent biological signatures were then transferred to several previously published data sets from fetal brain (Camp et al., 2015; Darmanis et al., 2015; Li et al., 2018; Zhong et al., 2018). This enabled the finer characterization of cell identities and the biological process they are involved in, which would not be feasible without transferred information. In addition, the resilience of overcorrection allows the detection of the possible contamination data source. Through integration across species, we identified shared sources of transcriptomic heterogeneity between mouse and human cells along the embryonic stage of interneuron development. The findings shed light on interneuron fate determination and maintenance across species. In aggregate, these analyses highlight the diversity of potential applications of the proposed method to borrow strength across multiple data sets and transfer to explore similar data.

## 2 Results

### 2.1 Methods description

cFIT models the scRNA-seq expression of individual cell using its cellular identity and domain-specific factors. Here *domain* refers to any standalone data set profiled at a single lab using a single technology from one batch (Figure 1A). Specifically, we are given a total number of *N* cells, and each cell *i* is associated with a p-dimensional feature vector *x_i_* corresponding to its gene expression values. Each cell comes from a specific domain with a known domain ID. Given *M* different domains, we use *m_i_* ∈ {1,2,…,*M*} to denote the domain ID of cell *i*.

**Figure 1:**
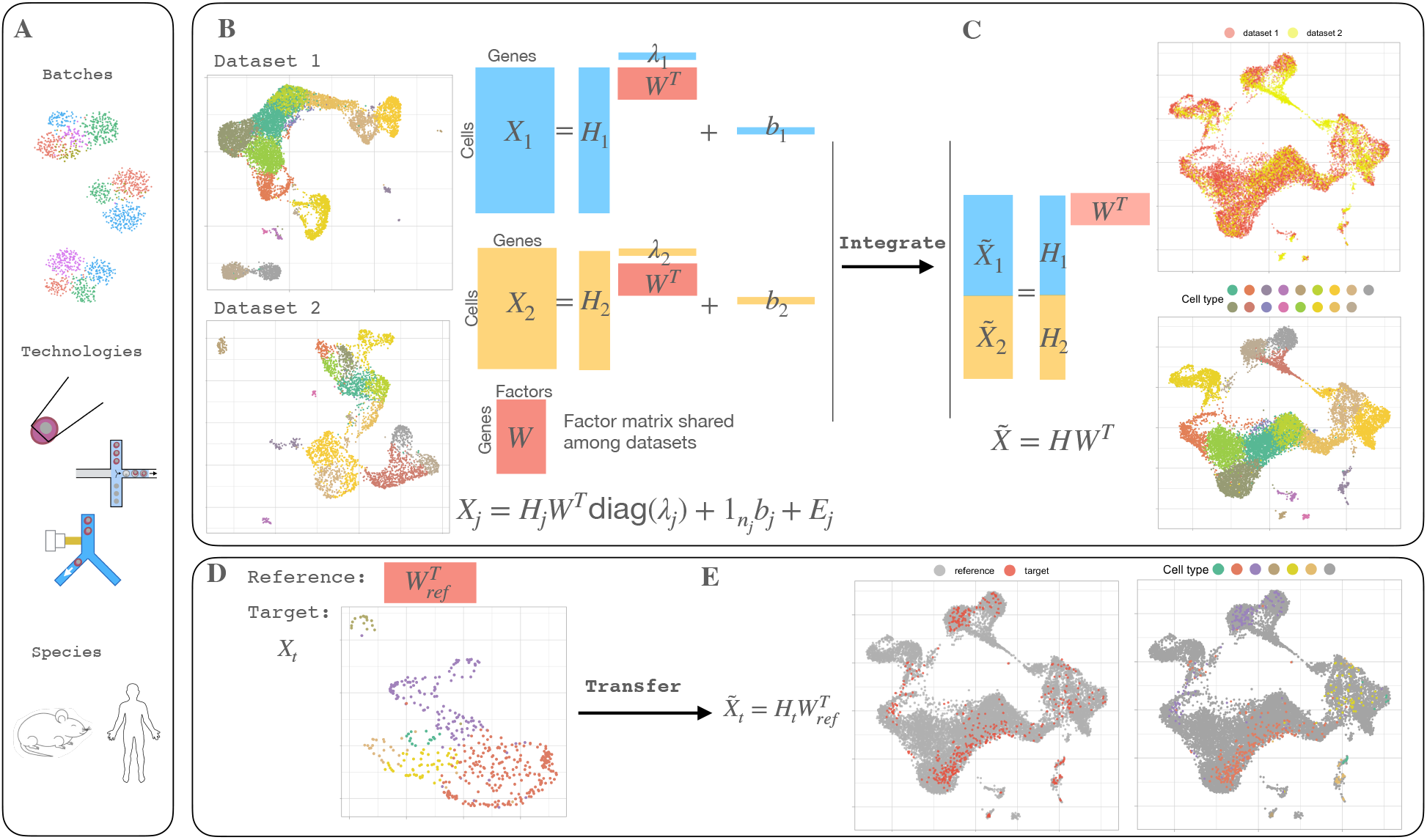
cFIT integration and transfer approach overview. (A) cFIT performs integration or transfer among scRNA-seq data sets from different batches, technologies, and across species. (B) Data integration takes in two or more data sets from different domains, where some cell-level biological processes are shared. Each data set is modeled by a low dimensional latent space corresponding to gene-level features (gene expression signatures), *W*, shared across domains, domain-specific factor loading *H_j_* characterizing cell composition, and domain-unique scaling, *λ_j_*, and shift, *b_j_*, capturing the technical distinction. (C) The integration algorithm estimates the set of parameters through iterative nonnegative matrix factorization (NMF). The integrated data can be obtained by eliminating the technical distinctions and projected onto a common subspace, where downstream analysis can be performed, such as clustering and trajectory inference. (D) The transfer process takes a reference factor matrix representing the gene-level signature profiles, and a new data set sharing the signature space. (E) The transfer algorithm estimates the target-specific parameters to project the target data onto the same low dimensional space as inferred from reference data. Cell labels can be assigned directly with unsupervised learning in low dimensional space or querying reference data.

We model the single cell RNA-seq data via a high-dimensional linear model with a latent low-dimensional structure. Let *x_i_* be the observation of cell *i* that is generated from

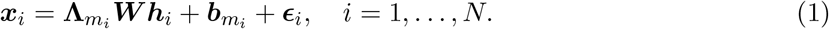

Here we use the diagonal (*p* × *p*) matrix **Λ**_*m_i_*_ = diag(**A**_*m_i_*_) to control the discrepancy of individual gene expression resulting from domain-specific technical effects. The matrix ***W*** denotes a (*p* × *r*)

non-negative factor matrix shared across all samples and domains representing the gene expression profiles (signatures) associated with the cells; each vector ***h_i_*** is a cell-specific non-negative vector of length *r* representing the factor loading vector of cell *i*; the domain-specific vector ***b**_m_i__* is a nonnegative vector of length *p* that captures the domain-associated shift. The noise terms 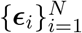 are modeled as independent, normally distributed random vectors with mean **0** and variance 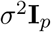, to account for measurement error from various sources.

Let *n_j_* denote the number of cells from batch *j* and 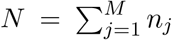 the total number of cells. Concatenating the scRNA expressions of all cells from each domain *j* as an (*n_j_* × *p*) matrix ***X**_j_*, then model (1) in matrix form is

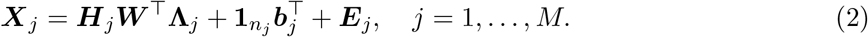

Here ***H**_j_* is a non-negative factor loading matrix, and ***E**_j_* is the noise matrix (Figure 1B).

Note that ***H**_j_* captures the biological heterogeneity originated from disparate cell type compositions, and {**Λ**_*j*_, ***b**_j_*,} are domain-specific parameters that accommodate domain differences such as batch effects from samples and libraries, different sequencing technologies, and even species; whereas ***W*** is the common factor space — the information shall be extracted and transferred to the new data set. The above model (cf. (2)) hinges upon the rationale that all the samples belong to, after proper shift and rescaling, the same lower dimensional linear subspace, which makes it possible to leverage information from diverse data sets.

To recover the set of unknown parameters 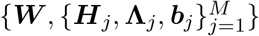, we propose to minimize the following objective

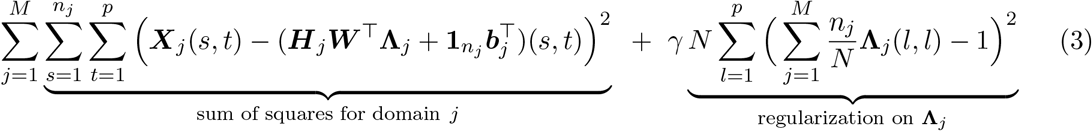

subjecting to the non-negative constraints for parameters ***W***, ***H**_j_*, **Λ**_*j*_, ***b**_j_*, where we use ***X***(*i,j*) to denote the (*i,j*)-the entry of matrix ***X***. Here, the positive parameter *γ* determines how much penalization is imposed on the batch-specific parameter **Λ**_*j*_, which ensures the model’s identifiability. To optimize the above non-convex objective function, we adopt the widely-used block coordinate descent approach (Bertsekas, 1976; Wright, 2015); see Section 4. Having eliminated the domain-related factors, the integrated scRNAseq expression matrices are ready for down-streaming analysis such as, clustering and trajectory inference with the low dimensional representation ***H**_j_*, and inference of differentially expressed genes with the scaled cell-by-gene matrix ***H**_j_* ***W***^⊤^ (Figure 1C). With the estimated common factor matrix ***W***, one can rapidly transfer the learned pattern to a new data set (target data) through the transfer algorithm. The data set-specific factor loading ***H*_target_**, scaling **Λ**_target_ and shift ***b***_target_ are estimated with fixed reference factor matrix through a similarly block coordinate descent procedure. Ultimately the algorithm projects the target data onto the low dimensional space estimated from the reference data set (Figure 1D). This procedure capitalizes on available information gleaned from prior analyses to optimize the value of small and low-quality data sets.

#### Comparisons to other methods

LIGER (Welch et al., 2019) is a popular linear method that employs a similar matrix factorization: it factorizes each batch expression matrix into a shared factor matrix *W*, similar to ours, for the joint embedding of cells across batches. Additionally, it introduces *p* × *r* parameters per data set to describe the domain effects. By contrast, we use only 2*p* parameters. By imposing a structural constraint on the data set-specific effect in cFIT, we restrict the domain-specific effects to the deviation by scale and shift, but rely on the shared factor matrix to model the relative signature matrix. The remaining orthogonal effects are preserved as biological distinctions. Though our model is comparatively conservative, we shall show in our data analyses that the structured model is sufficient to capture and remove the domain effects from various sources.

As comparisons, nonlinear methods typically involve the identification of mappings between data sets and remove the differences accordingly. One way of achieving this is by finding the mutual nearest neighbors (MNN), as used in MNNcorrect (Haghverdi et al., 2018), where the target data is mapped to the query data, guided by the pairwise points identified by MNN. A similar idea is employed in Seurat v3 (Stuart et al., 2019), which first uses canonical correlation analysis (CCA) to construct a shared subspace for two batches, and then identifies MNN across the two data sets (known as anchor points) to build locally weighted mappings. Apart from MNN-based methods, another type of nonlinear method leverages the alignment between clusters (distinct cell types) (DePasquale et al., 2019; Li et al., 2020) to eliminate the batch effects within the clusters and/or between mapped clusters. In the aforementioned methods, the success of MNN relies on the assumption that the batch effect is almost orthogonal to the biological subspace, and there is a substantial overlap of the cell compositions between the source and target data. As such, cluster-based methods require relatively well-separated clusters mapped across data sets. Apart from that, both types of approaches assume that the biological differences dominate the domain effects; however, this assumption is likely violated when cells are obtained along a continuous developmental trajectory with transitioning subtypes. These nonlinear methods eliminate the pairwise distinctions regardless of the source of distinctions. Therefore they tend to over-correct the biological variance. Alternatively, our model imposes no assumption on the population of cells in each individual data set. When our model assumptions are violated, cFIT simply fails to remove the batch effects rather than falsely erasing signals. In this sense, cFIT is less prone to over-correction.

In summary, by construction cFIT is resilient to over-correction of biological variance due to its simplicity, in comparison to existing widely-applied methods. We illustrate in practice that cFIT can successfully integrate data sets consisting of cells from various developmental stages along continuous trajectories and with very different sample compositions.

### 2.2 Simulation studies: differential expression analysis and robustness

The performance was evaluated in terms of differential expression gene (DEG) discovery, a key downstream analysis. The comparison assessed the integration methods’ ability to remove domain-specific factors while preserving biological signals. We compared cFIT with two widely-used singlecell integration methods: Seurat v3 (Stuart et al., 2019) and MNNCorrect (Haghverdi et al., 2018), which performed well in a benchmarking study (Tran et al., 2020). We do not compare with LIGER because this method does not produce a reconstructed scRNA expression matrix.

We simulated data under 5 settings with a combination of balanced/unbalanced batches, regular/high dropout rates, and two/multiple biological groups. After obtaining the reconstructed expression matrices, we used the two-sided Wilcox rank-sum test for DEG detection and reported the false discovery rate (FDR) among top 50 and 100 discoveries (Figure 2A). All three methods are effective in capturing the biological signals in some settings. Seurat v3 has advantages in the unbalanced batch/multiple group settings, while MNNCorrect shows better performances in the balanced batch scenarios. Overall, cFIT has the smallest FDR for most settings and is most robust across different parameter settings. More details can be found in Section 4.4.

**Figure 2:**
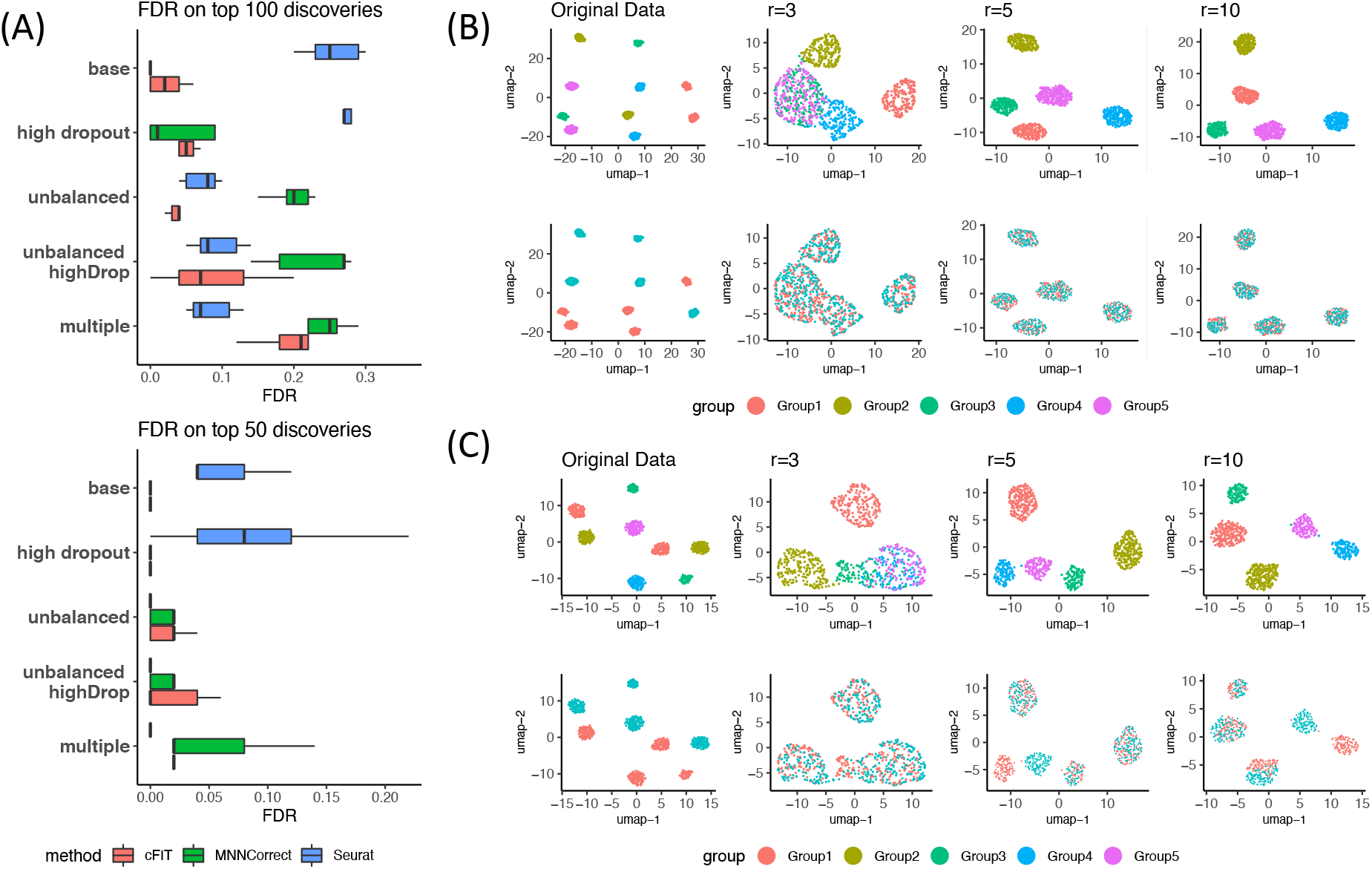
Evaluation and comparison using simulated data sets. (A) The False Discovery Rate (FDR) of the top 50 and 100 differential expression genes discovered by cFIT, Seurat v3, and MNNCorrect. Different combinations of parameters cover different scenarios of cell population sizes, drop-out rates, and the number of biological groups. Among the three addressed methods, cFIT has the smallest FDR for most settings and is most robust across different parameter settings. (B)(C) UMAP visualization of cFIT with different choices of *r*. The first row of plots are colored by cell group, and the second by batch. The latent dimension *r* should be equal to or larger than the actual number of biological groups. cFIT produces stable results when *r* is chosen from a reasonably wide range of values.

For the second part of our simulation, we evaluated the robustness with respect to tuning parameters. In Methods, we show that, with our proposed penalty term and parameter constraints, cFIT is identifiable and guaranteed to converge; furthermore, performance is enhanced using our initialization process. Hence the only key tuning parameter remaining is *r*, the number of latent factors. Here we demonstrate that our results are robust when *r* is chosen from a reasonably wide range.

By construction, the latent dimension *r* corresponds to the number of biological groups. We designed two simulation data sets such that the expression matrices follow model (2) with *r* = 5 and (i) all 5 cell groups presented in both domains; (ii) 3 cell groups presented in both domains, and each domain has a unique cell group. A UMAP visualization revealed the impact of severe batch effects (Figure 2B,C). Provided *r* ≥ 5, cFIT performed well, successfully integrating cells in each biological group into one cluster. Note that the second setting was challenging for most current integration methods, but cFIT was still able to recover the “unique cell groups”. More details can be found in Section 4.4. In practice, when we do not have prior knowledge about the number of biological groups, taking into consideration the complexity of real data sets and computation cost, we recommend a choice of *r* between 10 and 30.

### 2.3 Applications to single-cell data

#### Integration of scRNA-seq data sets from multiple technologies

Next, we evaluated our approach on human pancreatic islet cell data sets produced across five technologies, CelSeq, CelSeq2, Fluidigm C1, SMART-Seq2, and InDrop. Before correcting technical differences, cells were clustered by the data set of origin and cell type (Figure S1A). After applying the cFIT integration procedure, technical distinctions between data sets were effectively removed so that cells belonging to the same cell type, regardless of data sources, were well mixed (Figure S1B). With the integrated data, in addition to detecting all major cell classes (alpha, beta, delta, gamma, acinar, and stellar), we also identified some rare cells types that could not be reliably detected with a smaller number of cells through individual clustering analysis (schawann, mast). We benchmarked the performance of cFIT against two popular methods: LIGER (Welch et al., 2019) and Seurat v3 (Stuart et al., 2019). The integration results were compared using the alignment score, a measurement of how well different data sets mix (see Section 4), and the accuracy for preserving the cell type structure. Methods that perform well in both metrics effectively matched populations across data sets without blending distinct populations. Results show that cFIT achieved comparable high alignment scores and clustering accuracy as Seurat, indicating the effectiveness of our method relying on a simpler linear model compared to the nonlinear multi-step procedure employed by Seurat. LIGER over-corrected the domain-specific effects by falsely removing the biological gene-level distinctions between cells from different cell types. As a result, multiple major types were mixed together in the integrated data (beta, delta, gamma), resulting in lower clustering accuracy (Figure S1C,E). LIGER promoted the use of a post-quantile-normalization step to further align the quantiles of the batches within obtained clusters. Similarly, we show that this step can also be coupled with cFIT and produces higher alignment scores, despite the increased risk of mixing distinct cells types (Figure S2C).

To examine the robustness of the proposed method on integrating populations with different cell type compositions, we removed all cells of one pre-chosen type from each data set. The cell type was chosen as the largest or second-largest major cell type for each data set, while ensuring that the overlap of dropped cell types chosen was minimized (see Section 4). The integration procedure was applied to these perturbed data sets to compare performance. cFIT successfully integrated the data sets without blending distinct cell types. As with the original data, cFIT achieved high clustering accuracy (ARI ≈ 0.9) and alignment score (Figure 3A,B, Figure S2A,B). In contrast, LIGER and Seurat integration failed to preserve the cell type structure, and cells from different cell types were mixed together. Specifically, in both LIGER and Seurat results, a fraction of ductal cells were clustered with acinar cells, and beta, delta, alpha, gamma cells became entangled. The clustering accuracy (ARI) dropped to approximately 0.6 (Figure 3B). In summary, cFIT successfully characterized the domain-specific effects and was robust to the perturbation in relative cell type compositions.

**Figure 3:**
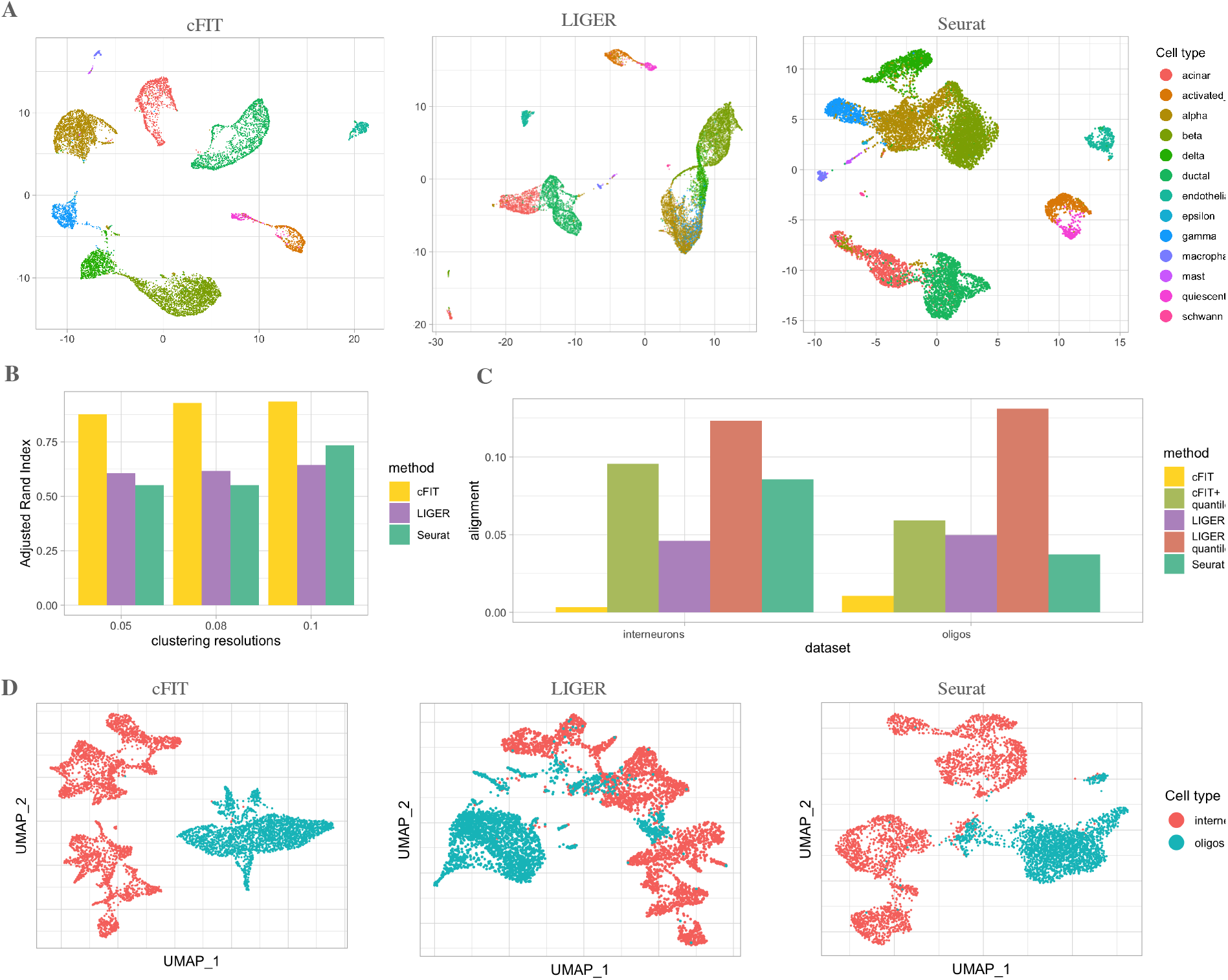
Integration of data sets with non-identical cell type compositions. (A) UMAP plots of integrated data from perturbed pancreas data sets, created by moving one major cell type from each data set, comparing results from three methods: cFIT (left), LIGER (middle), and Seurat (right). Cells are colored by cell type. (B) Evaluation of how well cell type structure is preserved from different integration methods applied on perturbed pancreatic islet data, though accuracy (ARI) of clustering on the integrated data set. A value close to 1 indicates better preservation of cellular transcriptomic identity. (C,D) Integration results of two data sets that are composed of distinct cell populations, oligodendrocytes, and interneurons correspondingly. (C) The alignment score was calculated on integrated data as a measurement of how well two data sets are aligned. (D) UMAP plot of the integrated oligodendrocyte and interneuron data sets from three different methods.

#### Integration of two data sets consisting of distinct cell types

To further ensure that cFIT was able to distinguish between the technical and biological effects when they were confounded, we jointly analyzed profiles of hippocampal oligodendrocytes and interneurons (Saunders et al., 2018). The two cell classes share a common origin in mouse development, but are born in distinct time frames and have very different functions in the mature brain. Therefore, we expected the two sets of mature cells share few, if any, common cell populations. Compared to LIGER and Seurat, cFIT generated the minimum false alignment, which is apparent visually in the UMAP display of integrated data and by comparing the alignment scores (Figure 3C,D). LIGER was most susceptible to over-correcting the internal substructure, which is particularly undesired for delineating distinct cell types.

#### Integration of human fetal cortical scRNA-seq data

Several studies, all following a unified set of protocols, have dissected multiple regions of fetal human brain and sequenced single cells (Camp et al., 2015; Darmanis et al., 2015; Nowakowski et al., 2017; Li et al., 2018; Zhong et al., 2018; Polioudakis et al., 2019). These studies enable the study of cellular programs in early development, comparisons of regional differences in cell type compositions and expression profiles, and mining associations between brain cell types and neurological disorders (Polioudakis et al., 2019; Satterstrom et al., 2020). However, challenges remain to integrate these data sets, which have been produced using disparate technologies, providing different coverage of developmental stage and brain regions. As a first step, we integrated the two largest scRNA-seq data sets from fetal cortex: (1) the Polioudakis scRNA-seq data (Polioudakis et al., 2019) (Drop-seq) of 33,976 cells from cortical anlage at gestation week 17 or 18 (GW); and (2), the Nowakowski data (Nowakowski et al., 2017) (Fluidigm C1) of 4,261 cells from primary cortical, medial ganglionic eminence (MGE), germinal zone, and cortical plate from primary visual cortex (V1) and prefrontal cortex (PFC). For the latter, samples ranged in age from 5.85 to 37 post-conception weeks (PCW). We aimed to borrow the advantages from both data sets via their integration, thus providing a more comprehensive characterization of human neocortical development. Directly concatenated, the normalized data exhibited substantial batch effects, especially among excitatory neurons (Figure 4A). By applying our proposed algorithm on these data sets, we obtained integrated data with good overlap in the UMAP two-dimensional space (Figure 4C). We observed several interesting results in the integrated data:

**Figure 4:**
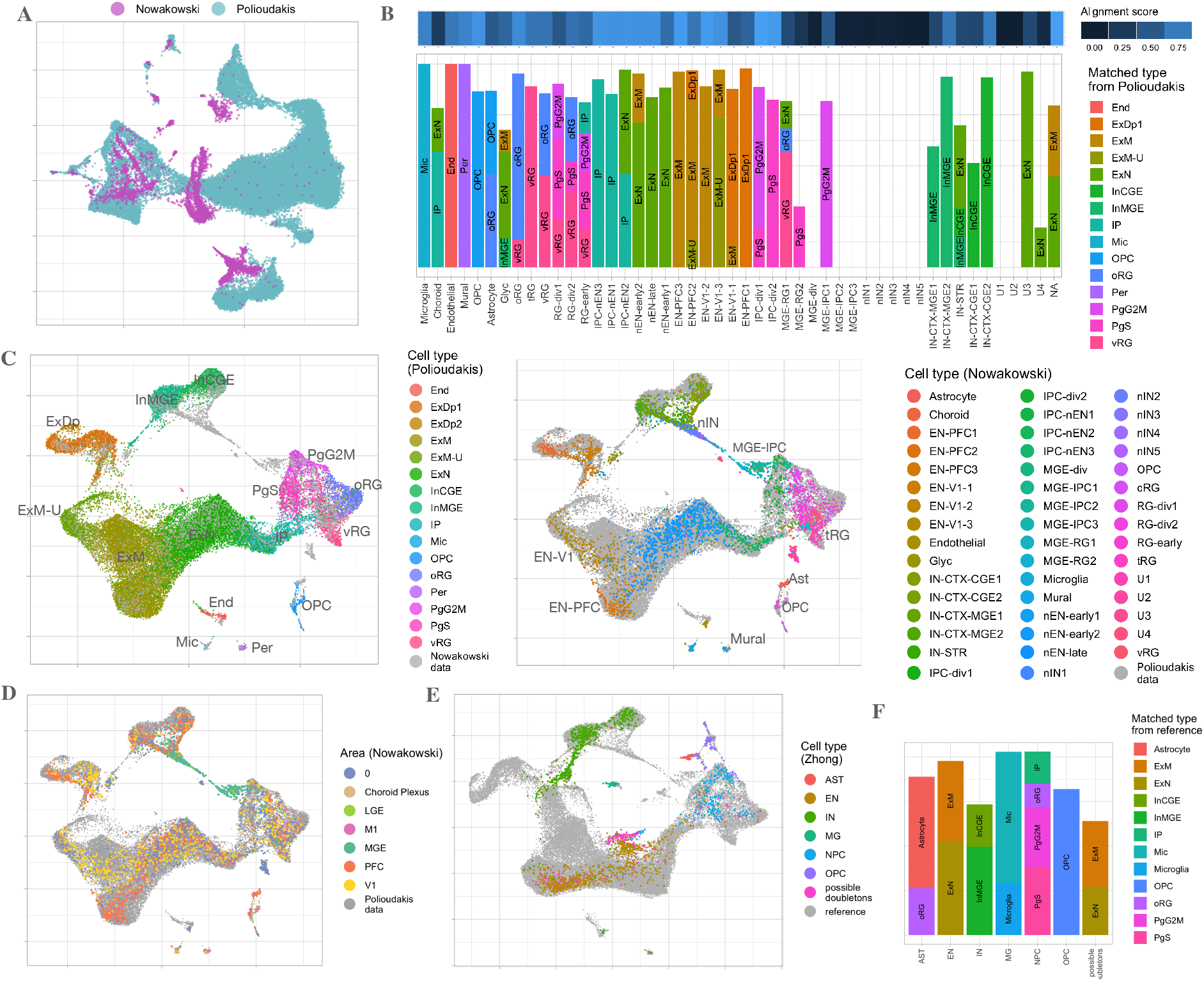
(A-D) Integration of two scRNA-seq data from the fetal brain (Nowakowski and Polioudakis data). (A) UMAP plot of two data sets before integration, but after depth normalization, log transformation, and per gene scaling (not centered). (B) Match of the identified clusters in Nowakowski data with major types identified from the Polioudakis data. The alignment score demonstrates how well each cluster in Nowakowski data is aligned with cells from Polioudakis data visualized in the color bar above. A higher score means the cells are well-matched with cells from the other data set. The bar plot shows the top matched cluster from Polioudakis data for each cluster of the Nowakowski data. (C) UMAP of scaled factor loadings obtained from data integration. In left panels, only cells from Polioudakis data are colored into 16 major cell types. Similarly, in the right panel, only cells from Nowakowski data are colored according to the 48-cell-type label. (D) UMAP of integrated Polioudakis and Nowakowski data, colored by brain area annotated for Nowakowski data. (E-F) Transfer results on 2309 cells from Zhong data. (E) UMAP of 2309 cells (colored by Zhong labels) overlaying on cells from Polioudakis and Nowakowski (reference) data sets, among them a group of cells outside the range of reference cells and previously identified as possible doubletons (Zhu et al., 2019). (F) The composition of cells based on matched cell types in reference data sets for each major group, with the alignment score computed for each major group measuring how well it matches with reference data.

First, the two data sets align substantially, as indicated by the alignment score (Figure 4B). Some similarly labeled cell types matched perfectly, including OPC, radial glia, progenitor cells, excitatory neurons, interneurons, and some rare types like microglia (Figure 4B). Others were unique to Nowakowski data, including interneuron progenitors (MGE-RG, MGE-IPC, nIN) and unknown types (U1, U2). The interneuron progenitors were sampled from a brain region not included in the Polioudakis data set; the unknown types could be other unrepresented cell types or artifacts. It was notable that cFIT did not force these cells to overlap with closely related cell types present in the larger Polioudakis data set.

Second, two maturation trajectories of excitatory neurons were revealed in the Nowakowski data. One ends at PFC maturing excitatory cells, and the other ends at V1 maturing excitatory cells; while the mature cells sampled from the two different brain regions were clearly differentiated, the immature cells were not (Figure 4C,D). The Polioudakis data spanned this same space, but this study did not distinguish cells by region. Thus, by aligning the two data sets and noting the two trajectories apparent in the Nowakowski data, we can infer the regional origin of a portion of the Polioudakis cells (Figure 4D). In the Nowakowski data, there was a continuous trajectory from the progenitor cells to interneurons, labeled in the paper as the MGE progenitors, newborn MGE neurons, MGE-interneurons, and CGE-interneurons. This trajectory was not identified in the Polioudakis data set because it does not include samples from the MGE/CGE/LGE region, which is precisely where one would expect to find interneuron progenitor cells.

Third, we were able to label the previously unannotated cells (NA) in the Nowakowski data as excitatory neurons via their substantial overlap with migrating and maturing excitatory neurons in the Polioudakis data and the resulting high alignment score. Some cells in two unknown cell clusters (U3, U4) can also be assigned as newborn excitatory neurons because they overlap with ExN labeled cells in the Polioudakis data. In addition, we identified a subset of 39 OPC cells in Polioudakis data that are likely astrocytes, given they are matched to astrocytes from the Nowakowski data and expressed astrocytes markers GFAP, SOX9, and EGFR.

#### Transfer the learned signature from a data set with a larger number of human cortical cells to a data set with a smaller number of human cortical cells

The learned factor matrix from the integration of the Nowakowski and Polioudakis data can serve as a comprehensive reference that characterizes the cellular processes in the fetal brain, covering a wide range of developmental stages (6-37 weeks) and major and more specialized cell types. Thus, we applied our proposed transfer learning methods and using this comprehensive reference to the Zhong data set (Zhong et al., 2018), which contains 2309 single cells from the human embryonic prefrontal cortex (PFC) from 8 PCW to 26 PCW. Using the Seurat package (Stuart et al., 2019), the authors identified six major clusters: neural progenitor cells (NPC), excitatory neurons (EN), interneurons (IN), astrocytes (AST), oligodendrocyte progenitor cells (OPC) and microglia (MIC); these will be referred to as Zhong labels. Through the transfer procedure, we obtained the following results: First, the 2309 cells, represented in low dimensional space using estimated factor loadings, overlay substantially with cells from the Polioudakis and Nowakowski data sets. We were able to identify finer structure within each major group by matching group labels from the reference data sets (Figure 4E). Among them, NPC consisted of cycling progenitors, intermediate progenitors, and radial glia; EN cells contain both migrating and maturing excitatory neurons; IN matched perfectly with MGE and CGE interneurons; OPC and MIC aligned perfectly with OPC and microglia from reference data; and AST aligned partially with astrocytes and partially with oRG. Most groups showed high alignment scores, except for the astrocyte (AST) clusters, which while laying adjacent to each other, were not well mixed. This analysis showed that by transferring rich information, we were capable of delineating finer structures within each major type in another data set. Second, we were able to identify cells from rare types that were likely mis-clustered in the Zhong analysis. For instance, previously labeled as interneurons (IN), five cells were relabeled as endothelial, and another five cells were relabeled as pericytes. This finding was supported by the UMAP visualization, where these cells lie in clusters corresponding to endothelial and pericytes that were well separated from other types. Third, the migrating and maturing excitatory neurons align well with the PFC branch as identified from the Nowakowski data (the branch extends downward in the UMAP visualization Figure 4C), while the upper branch displays maturing excitatory neurons from the visual cortex. Fourth, we observed a fraction of EN cells that align poorly with the reference data (outside the gray range in UMAP plot), among them some were labeled as NPC. This was indicative of the under-representation of these cells in the reference. It was not clear whether these were novel neuronal signatures characteristic of these cells or were mischaracterized cells. The latter is more likely because previous work (Zhu et al., 2019) discovered a cluster of cells (marked in Figure 4) as possible doubletons (i.e., transcripts captured from two cells rather than one).

Transfer learning is particularly valuable when applied to much smaller data sets. The Li data (Li et al., 2018) contains 762 cells collected from nine brains ranging in age from 5 to 20 PCW. These cells derive from the fronto-parietal neo-cortical wall, mid-fetal fronto-parietal neo-cortical plate, or the adjacent subplate zone. Applying cFIT, the cells were transferred to the reference data in the low dimensional factor space and showed good alignment of most cell labels (Figure S3A,C). The alignment was further validated by the incremental age along the trajectory observed (Figure S3B). By transferring onto well-characterized developmental trajectories, we were also able to identify some likely mislabeled cells, which likely arose due to having too few cells per type available originally: for instance, the rainbow points along the maturation trajectory of excitatory neurons and CGE/MGE interneurons. We next examined an even smaller scRNA-seq data set from the fetal brain, with 220 cells sampled between 12 and 13 PCW (Camp et al., 2015). Guided by marker genes, these single cells were previously labeled into seven types in the original paper: two subtypes of apical progenitors (AP1, AP2), two subtypes of basal progenitors (BP1, BP2), and three subtypes of neurons (N1, N2, N3). By transferring these cells onto the reference data, we were able to validate the cell labels (Figure S3D). As a further step, we identified the finer structure within each cell group (different types of radial glia within AP cells and different progenitor cells within the BP group; Figure S3E). We also revisited a widely studied data set composed of both fetal and adult brain cells (Darmanis et al., 2015)(134 fetal cells and 334 adult cells from cortical tissues at 16-18 PCW). The transfer procedure evenly distributed the fetal cells along the reference trajectory, where labels were easily inferred (Figure S3F). By contrast, the transfer process projected all adult neurons outside of the fetal developing trajectory (Figure S3G).

#### Integrate cell expressions across species

We further examined the performance of the proposed algorithm for integrating cells across species, specifically mouse and human. Mouse models have proven useful for understanding the basis of many human diseases, in part because of the ready access to tissue from which to generate data. Although we expect differences between the two species, similarities have also been noted in some transcriptomic patterns (Wang et al., 2018) and, once understood, shared features are likely to provide a deeper insight into the fundamental architecture underlying cellular development and physiology.

Diverse subsets of cortical interneurons have vital roles in higher-order brain functions. To investigate the molecular diversification of early inhibitory precursors, we leveraged several mouse scRNA-seq data sets collected along a developmental time course using multiple technologies (Dropseq, 10X, SmartSeq2) (Mayer et al., 2018; Tasic et al., 2016) and integrated them with human medial ganglionic eminences (MGE) progenitors and interneurons (Nowakowski et al., 2017). We first examined the heterogeneity within the early development in MGE (mainly composed of mitotic progenitors within MGE before migrating to cortex). By integrating the 733 Nowakowski cells collected from MGE (age between PCW5 to 21) with 5622 mouse cells from Dropseq data (embryonic day (E)13.5), we identified a common mitotic developmental trajectory shared between the two species (Figure 5A,B). In their prior work (Mayer et al., 2018; Tasic et al., 2016), each mouse cell was assigned a maturation score (a continuous value quantifying the extent of cell development), allowing the cells to be divided into mitotic and post-mitotic stages. We mapped this maturation score to human cells by averaging over the scores of neighboring mouse cells (among the 30 nearest neighbors). We observed a match between the maturation score and the reference labels of human cells, starting from the two subtypes of MGE radial glia (MGE-RG), followed by the dividing MGE progenitors (MGE-div), subtypes of MGE intermediate progenitor cells (MGE-IPC), and ending with newborn interneurons (Figure 5C). Seurat clustering analysis was performed, which identified eight major clusters, each composed of both mouse and human cells (Figure 5D). Clusters A-F are composed of mitotic cells concordant with the reference labels (Figure 5E). Meanwhile, the two clusters of post-mitotic cells aligned with the reference branch labels. The newborn interneurons fell in branch 1 cluster, which was conjectured to give rise of cortical interneurons (Mayer et al., 2018). The alignment score calculated per cluster demonstrates that the human cells were evenly distributed in each cluster, except for F (Figure 5E). Cluster F contained a fraction of newborn interneurons from a later developmental stage (>20PCW), beyond the range covered by mouse cells.

**Figure 5:**
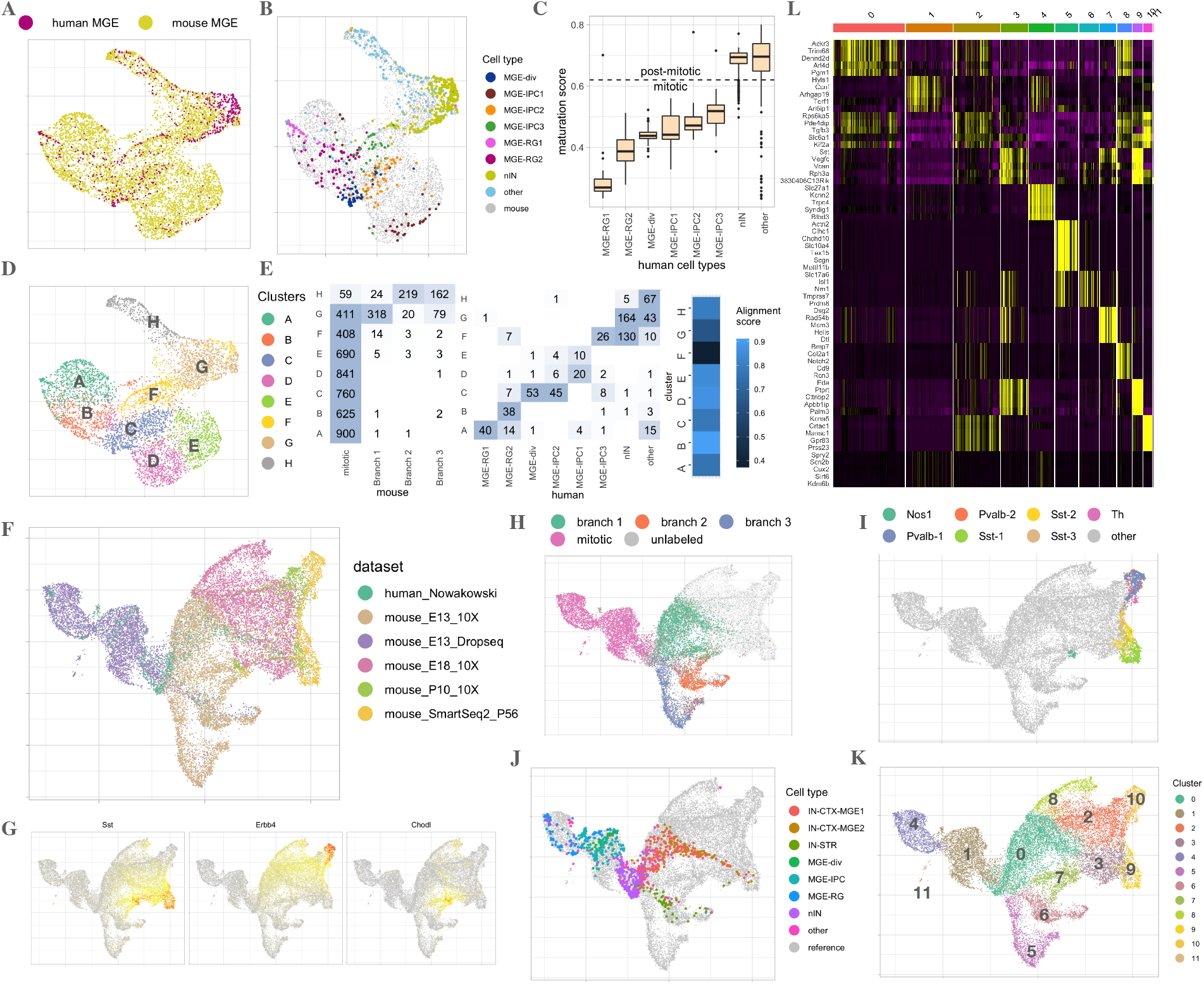
Integration of cells from medial ganglionic eminences (MGE) and cortical interneurons from human and mouse. (A-E) Integration of mouse cells from MGE (Dropseq) with human MGE progenitor cells. (A-B) UMAP plot of integrated data colored by data set (A), and colored by known human cell types (B). (C) mapped maturation score for human cells for each human MGE cell type. (D) Clustering analysis on integrated data identified 8 groups, ordered from A to H based on maturation scores. (E) Confusion matrix comparing the identified cell clusters with the reference mouse and human cell labels, with the alignment scores calculated to measure how well the human cells are blended in mouse cells in each cluster. (F-L) Full integration of six data sets, including mouse MGE cells (E13.5), interneurons precursors from mouse embryos (E13.5, E18), postnatal (P10), adult stage (P56), and human MGE and cortical interneurons. (F) UMAP plot of integrated data colored by the data set. (G) Expression of genes marking Sst interneurons, Pvalb interneurons (Erbb4), and Nos1 cells (Chodl). (H-J) Highlight among integrated data the mouse cells from three developmental branches (H), adult mouse cells (two subtypes of Pvalb, three subtypes of Sst, Nos1 and Th) (I), and human cells by reference cell types (J). (K) Integrated data were clustered into 12 distinct cell groups based on transcriptional differences. (L) Heatmap visualizing the top five marker genes for each identified cell clusters.

Next, we investigated the full developmental process starting from mitotic progenitors in MGE, which differentiate and migrate to the cortex to become mouse mature interneurons, and examined whether the human MGE-derived interneurons developmental trajectory could be aligned. We integrated six data sets, a human data set containing 733 cells from MGE and 271 MGE-derived cortical interneurons (Nowakowski et al., 2017) (pcw5-22), and five from mouse at different ages and sequenced by different technologies (Mayer et al., 2018), including 5622 mitotic and postmitotic cells from MGE (E13.5, DropSeq), 6515 post-mitotic interneurons precursors from mouse embryos (E13.5, E18 from 10X), 1269 cells at postnatal 10 days (P10 from 10X) and 1350 cells adult stage (P56 from SmartSeq2) (Tasic et al., 2016). We observed a continuous developmental trajectory from the integrated data (Figure 5F) starting from the mitotic cells, followed by those transitioning into the post-mitotic stage where progenitors diverge and differentiated by distinct transcriptional states. The states aligned with the previously identified three branches (Figure 5H). Along branch 1, new sub-branches emerged and ultimately arrived at distinct types of mature interneurons (Sst, Pvalb, Nos1, and Th; Figure 5I). The human cell development largely resembled the mouse cells, where the human cells ranged from early mitotic to relatively mature interneurons along branch 1. In another trajectory endpoint, striatal (IN-STR) interneurons were blended with mouse cells from branch 2(Figure 5J). This analysis identified 12 major groups, including three clusters of mitotic cells (11,4,1) and branch 1 clusters of four different sub-branches: an Sst branch (cluster 3,9); a Pvalb (cluster 2, 10); a Nos1 branch (cluster 7); and an unknown branch (cluster 8, expressing markers such as BmT7, Col2a1, Notch2, Cd9) (Figure 5K,L). The human cells were grouped with mouse cells in corresponding clusters (Figure S4B). Particularly, a group of human cells in cluster 3 were identified as relatively mature precursors of Sst interneurons (Figure S4B).

Our analyses reveal that transcriptional profiles underlying interneuronal fate specification are largely conserved between mouse and human. It also highlights the power of cFIT to capture and integrate the shared biological processes across different systems effectively. These are important steps towards the goal of harnessing information across species to understand mammalian neurodevelopment and its relevant physiology.

## 3 Discussion

As the field matures, a diversity of scRNA-seq data sets has been collected, which if integrated could provide valuable biological insights. The challenge is that these data vary in features, both large and small, ranging from different batches, different technologies, different developmental periods, different species, and even different modalities. We develop a method for integrating single-cell data data sets and transferring knowledge to new settings that is robust and interpretable. cFIT assumes a shared common factor space across data sets, but it models distortions and shifts on gene-wise expression that are unique to each source. In doing so, it captures the advantages without the disadvantages of existing methods. Like LIGER, the nonnegativity constraint of NMF yields interpretable factors that can be biologically meaningful. While the shift in cFIT corresponds to the batch-specific factor in LIGER, their scaling is applied to all the shared factors concurrently. There-fore, our model takes a different approach to characterize the impact of domain effects originated from different experimental tools and measurements employed. Seurat v3 intrinsically assumes a substantial overlap of cell-type compositions between target and source data sets, and produces results dependent on the order of pairwise integration. By contrast, our method makes no assumptions on cell populations in individual data and can perform integration simultaneously across all data sources. Our method also does not depend on the degree of domain-effects relative to biological differences (e.g., between cell types), as required to ensure the success of methods such as DESC (Li et al., 2020) and MNNcorrect (Haghverdi et al., 2018). Besides, far fewer parameters are required by our model, while retaining the power to capture domain effects. It maintains identifiability and robustness through the choice of tuning parameters. Like Seurat v3, our method also provides an estimate of a corrected expression matrix, which can be used as input for downstream analyses such as pseudotime or differential gene expression analysis. There are notable advantages to having access to both corrected gene expression and the noise reduction of visualizing lower-dimensional factor loadings that reveal interesting biological features.

Unlike many competing approaches, cFIT is less prone to overly correcting biological heterogeneity, which facilitates combining data sets with biological heterogeneity and capturing the advantages of each source into a single corrected data set. We show this in simulations and several scRNA-seq data sets. Consequentially, in our analysis of fetal brain development, we were able to combine data sets sampled from widely divergent protocols, spanning different developmental epochs of both mouse and human cells. The resulting integrated analysis delineated closely related neuronal subtypes, drew inferences about developmental trajectories, and correctly classified rare cells from data sets with small numbers of cells sampled. These insights could not be obtained from any single data set, nor with an integration approach that minimized biological differences between data sets.

Although our simulations show that a good initialization often leads to a close approximation of the global optimum, it would be satisfying to obtain theoretical justification for this observation. Given the non-convex nature of the optimization scheme, this is a challenging problem; however, recent work (Tripuraneni et al., 2020) characterizes the error behavior for estimating a shared factor space for a linear regression model. It would be interesting to see whether their ideas can be carried through in our setting.

We note that one promising area for future work that we did not yet explore is the integration of multi-omic data. Within a modality, cFIT overcomes the challenge of reconciling the considerable heterogeneity observed across individual data sets, but the variation across modalities such as single-cell ATAC-seq and DNA methylation can be substantially greater. Given the fundamental similarities between cFIT and LIGER due to the common factor approach, however, it seems likely that cFIT could be successfully extended to multi-omic data.

## 4 Methods

### 4.1 cFIT framework

In this section, we describe the cFIT method in detail, including how the unknown parameters are learned in a data-dependent manner, and how to transfer information to new data sets for more accurate decision-making.

As discussed in Section 2.1, cFIT learns the unknown parameters by minimizing the objective function (3). In a more compact form, we aim to minimize

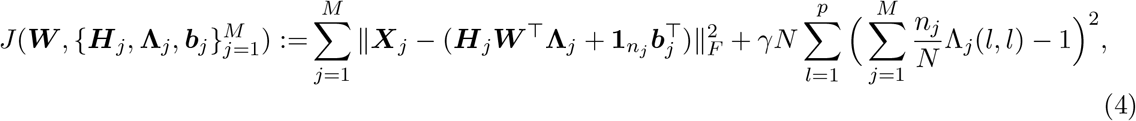

subjecting to the non-negative constraints ***W***, ***H**_j_*, **Λ***_j_*, ***b**_j_* ≥ 0. (Note that we write ***W*** ≥ 0 when each element of the matrix or vector is non-negative.) Here, the squared Frobenius norm of a matrix is defined as the sum of squares of all its entries. We remark that we impose the restriction that ***H**_j_* **1**_*r*_ = **1**_*n_j_*_ — which is also known as *row-stochastic assumption* (Fu et al., 2019) — to guarantee the model identifiability, since ***H**_j_* ***W***^⊤^**Λ**_*j*_ are in product form, one can always rescale ***H**_j_* and ***W*** otherwise. Additionally, the penalty term 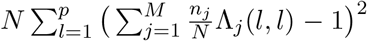 is introduced to ensure identifiability of the solution as ***W***^⊤^ ***L***^-1^ ***L*****Λ***_j_* = ***W***^⊤^**Λ**_*j*_ for any full-rank *p* × *p* diagonal matrix ***L***. (Theoretically 0 can always be achieved. In practice it is sufficient to fix the penalization parameter *γ* to a large value, e.g. *γ* = 10^6^.)

To summarize, our goal is to find the solution to the following optimization problem

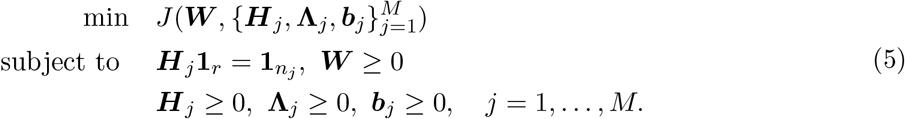

In comparison with LIGER (Welch et al., 2019), cFIT contains as a component similar domain-specific factors in the form of shifts 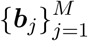, while it introduces an additional group of parameters 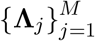 to characterize the distortion of gene-wise expression. The domain-specific scaling is applied concurrently across all the shared factors of each gene, capturing the discrepancy originated from different quantification and normalization tools employed under diverse experimental conditions. Therefore our approach is fundamentally different from LIGER that models the differences through higher-dimensional domain-specific factors. We provide the details of the optimization algorithm in the section below.

#### Data integration

Given the solution 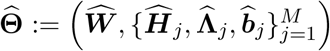, one can obtain the estimated data matrix in the shared lower dimensional factor space

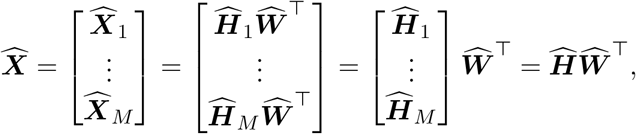

and the intrinsic low dimensional representation as the estimated factor loadings 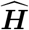. As is seen in our real data examples, these batch-adjusted observations are particularly useful for down-streaming analysis, such as cell-type identification.

#### Data transfer

Equipped with solution to the above optimization problem (5), we are able to transfer the knowledge to a new data set that potentially contains a fewer number of cells or is of lower quality. Toward this, we generalize the proposed model and algorithm for transfer learning. Given target expression matrix *X*_target_ and factor matrix ***W***_ref_ from reference data sets, the goal is to recover (***H***_target_, **Λ**_target_, ***b***_target_). We propose the folllowing objective

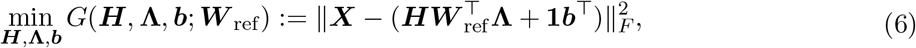

subject to the non-negative constraints ***H***, ***λ***, ***b*** ≥ 0 and row stochastic constraint ***H*1** = **1** (Algorithm 3).

### 4.2 Algorithm details

Notably, the objective function is non-convex with respect to the parameter space Θ whose optimization landscape is typically unclear. Nevertheless, the algorithmic properties and theoretical behaviors have been extensively studied in the literature of non-negative matrix factorization (NMF) (Lee and Seung, 2001; Donoho and Stodden, 2004; Wang and Zhang, 2012; Vavasis, 2010). Here we adopt the block coordinate descent approach with random permutations (Bertsekas, 1976; Wright, 2015). The algorithm proceeds in an iterative manner: at each round, randomly select a block of variables and take a gradient step with respect to this block while holding other blocks fixed; and then proceed iteratively until the algorithm converges.

Block coordinate descent is particularly useful in practice, as it turns an optimization problem of much higher dimensions into solving several sub-problems. In addition, it transforms the non-convex objective into several convex sub-problems, which can be carried out much efficiently. For convex objectives, it is known that block coordinate descent with random permutations converges to the optimal solution under very mild conditions (Wright, 2015; Sun et al., 2020).

In our case, it is easily seen that each sub-problem is convex, and the exact solution can be obtained by non-negative least squares (NNLS) with constraints. We solve the sub-problems alternatively in a randomly permuted manner to ensure convergence to *stationary point* (Sun et al., 2020) (Algorithm 1). Since the algorithm is not guaranteed to find the global optimum, parameter initialization can be critical. Intuitively, the binary membership matrix, achieved from an initial clustering of centered and scaled expression data, is suitable as a descent approximation of true factor loading matrix. Therefore, we initialize ***H**_j_* from clustering the concatenated and column-standardized expression matrices, and initialize ***W*** using NNLS after setting ***b**_j_* = **0** and **λ**_*j*_ = **1**_*p*_. It is shown empirically that the proposed parameter initialization provides a good starting point that speeds up the algorithm convergence (Algorithm 2).

#### Data pre-processing

To integrate multiple data sets, we first perform a feature selection step on each data set individually. Then prioritize those features which are often independently identified as highly variable across multiple selection sets. We select 2000 to 4000 (depending on sample size) top-ranked highly variable genes (HVG) shared among data sets, with ties determined by examining the ranks of the tied features in each original data set and taking those with the highest median rank (Stuart et al., 2019). After the common genes are selected, the expression of each cell is normalized by library size and log transformed (log(*x*/ ∑ *x*) * 10^4^ + 1). Though our method does not require scaling per gene in the pre-processing step, it is shown in empirical analysis that scaling each gene per data set helps to identify the rare types. It should be noted that we do not center the gene expressions since the algorithm takes as input the non-negative expression matrices. To transfer the learned factors to a new data set (target data), the rows of factor matrix *W*_ref_ is a subset of the genes that are also expressed in the target data. The cell expressions on the remaining genes in the target data are then normalized following standard pre-processing procedure (normalized by library size of a cell, log-transformed, and scaled per gene).

##### Algorithm 1: Data integration

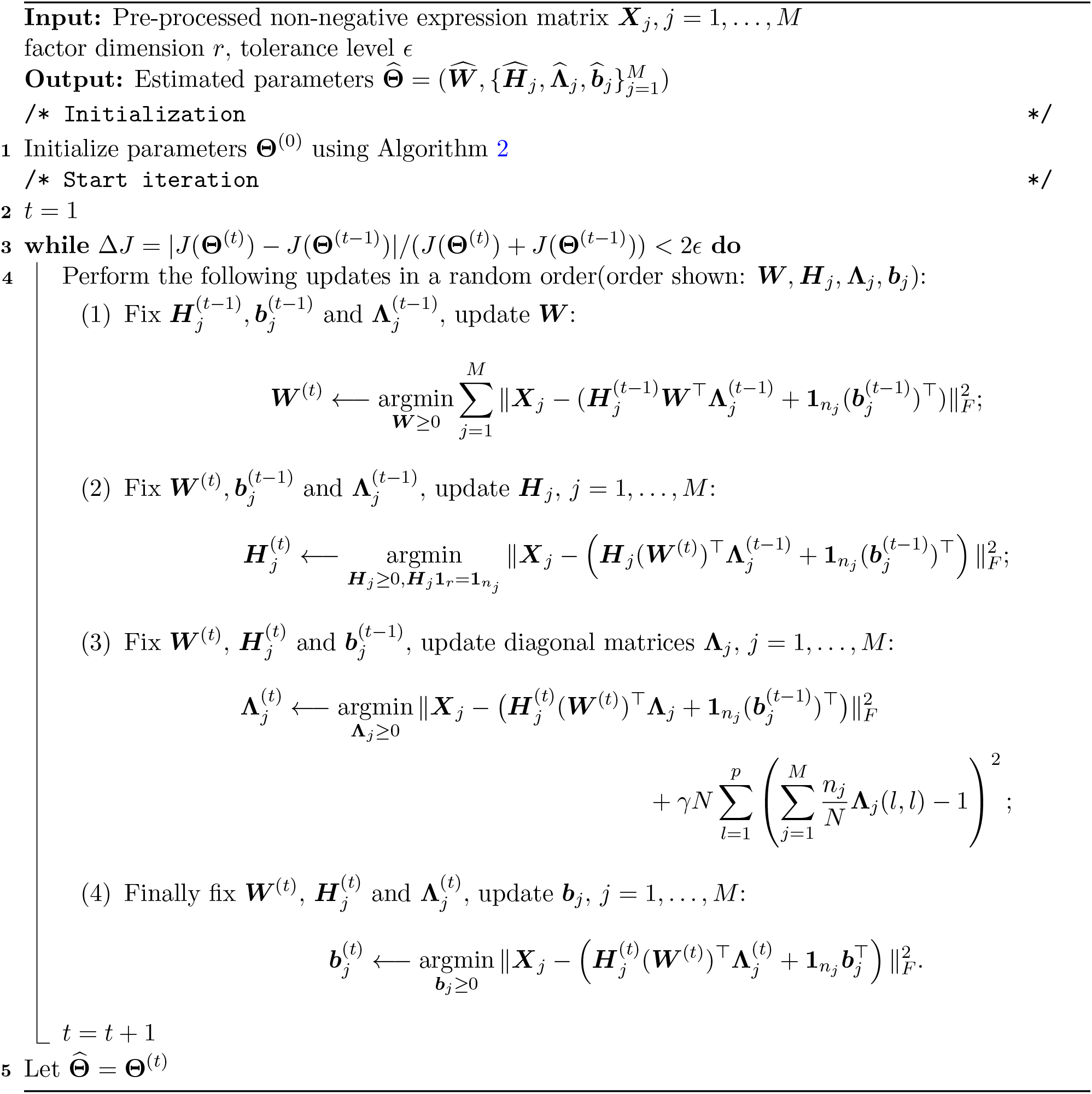

##### Algorithm 2: Parameter initialization

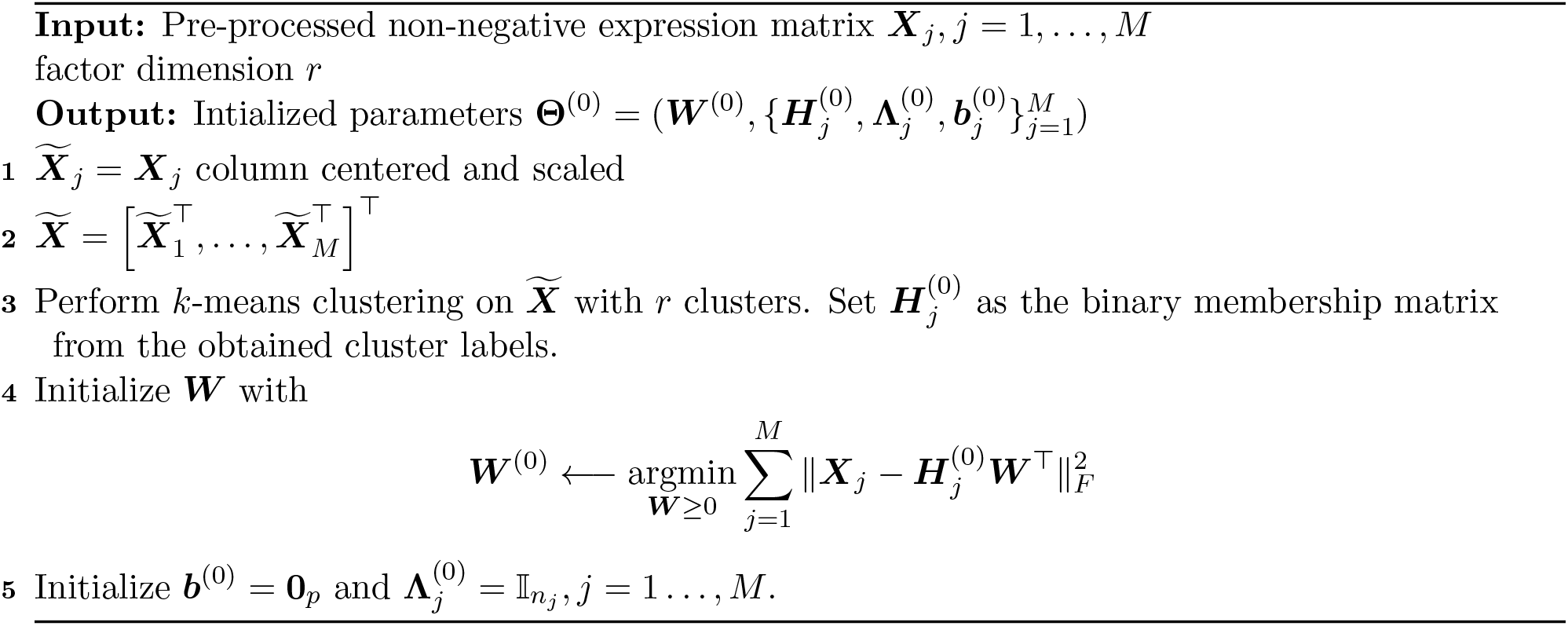

##### Algorithm 3: Data transfer

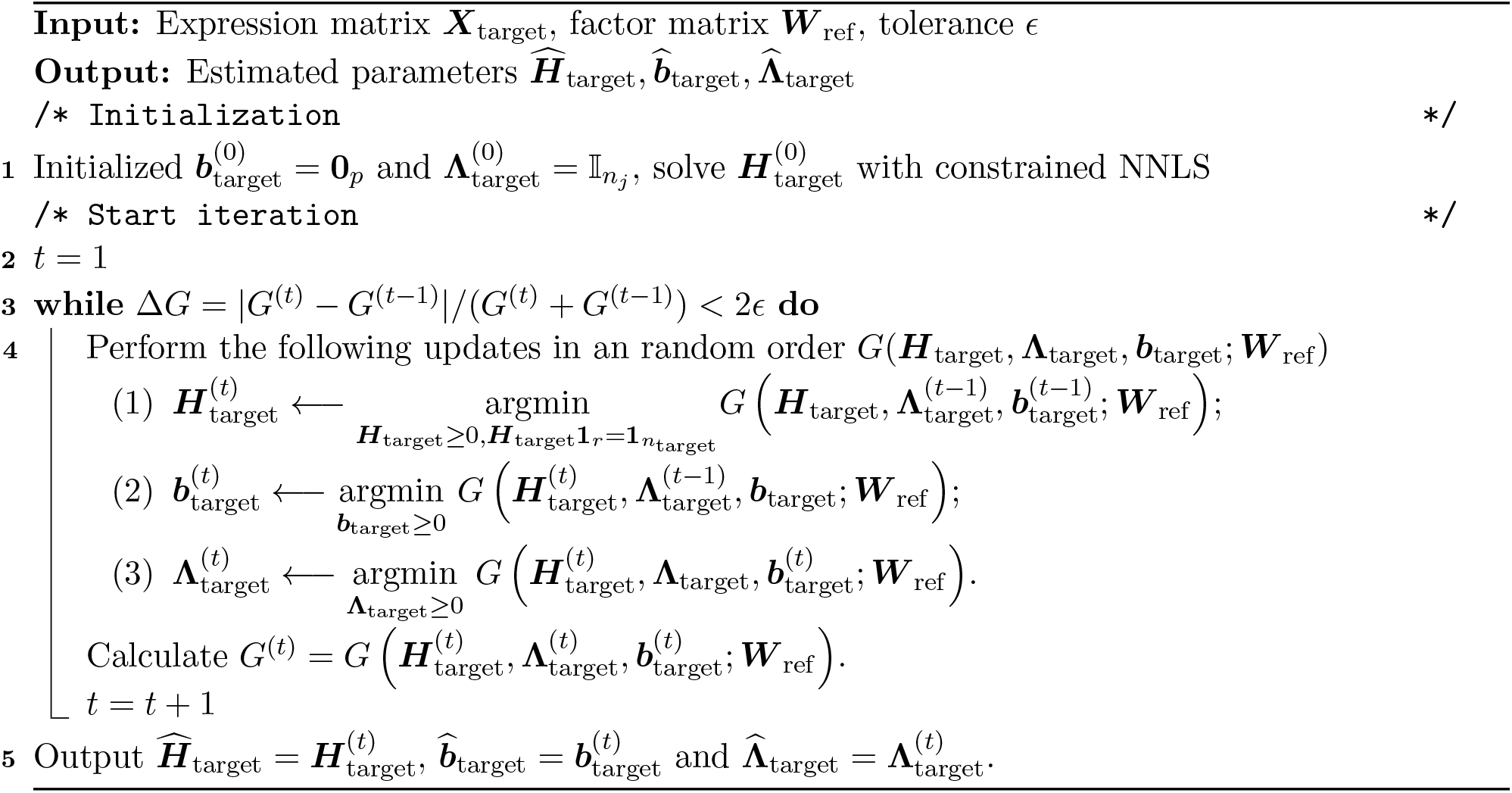

#### Integration post-processing

The factor loading matrices obtained via Algorithm 1 are post-processed, where each column is multiplied with the column sum of the factor matrix. The resulting low dimensional representations are ready for downstream analysis such as clustering and trajectory inference. To identify differentially expressed genes, the gene expression across cells in the integrated data can be recovered via ***HW***^⊤^, where each column corresponds to the gene-wise expression. We also examined the role of quantile normalization in the post-precessing step as proposed in LIGER (Welch et al., 2019). Specifically, with the estimated factor loading matrix, a reference data set is chosen (by default the data set with the largest number of cells), and the quantiles of the factor loadings for each joint cluster in the other data sets are normalized to match the quantiles of the reference data set for that joint cluster. This step is helpful to obtain more balanced and robust factor loading for smaller data sets.

### 4.3 Alignment score

The alignment score quantifies how well-aligned the two or more data sets are. We calculate the alignment metric as in Seurat v2 (Butler et al., 2018): First construct a nearest-neighbor graph based on the cell embeddings in low dimensional space using, for instance, the factor loadings provided by cFIT; Next, for each cell, compute the number of its *k* nearest neighbors that belong to the same data set as the alignment score for that cell; To obtain the alignment score for each cell type or each data set, the per-cell score is averaged over cells in the data set/cell type, denoted as 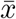. Finally, we normalize by the expected number of same data set cells and scale to range from 0 to 1, where a value of 1 represents perfect alignment. Mathematically, the rescaled alignment score can be written as

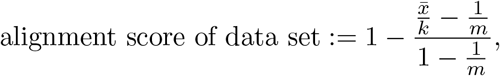

assuming data sets are of equal size and similar cell type compositions. When the size of data sets are widely different, instead of subsampling all the data sets down to the smallest, we address the disparate size by replacing the expected value 1/*m* with a data set-specific expected value 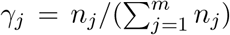 given the size of data set *j* being *n_j_*. It worth mentioning that in our real data examples, we seldomly observe that all source data sets follow the same cell type distribution. Therefore we use the alignment score mainly as a relative measure in our analysis (when comparing the performance of multiple methods applied on the same set of source data, or comparing the alignment of different cell types when the fractions of cells from each data set is comparable among cell types).

### 4.4 Simulation details

In this part, we describe the simulation settings. The Splatter simulation package(Zappia et al., 2017) permits us to control biological and technical factors, including library sizes, dropout rates, number of cell groups, and all the simulated data sets we describe below are generated with Splatter.

In the first simulation part, we simulated 10000 genes with differential expression probability set as 0.05. We implemented all the three methods on the same set of 2000 high variable genes selected by Seurat v3, among which the number of DEGs well exceed 100. The following table summarizes the batch size, group probability and dropout rates we used. The error bar in Figure 2A was obtained by running each scenario with 10 different random seeds. The dropouts were simulated by setting `drop.mid` as −4 and −2 in Splatter. We set *r* = 10 and maximal iteration number as 200 for the cFIT part, and used the default settings of Seurat v3 and MNNCorrect.

In the second simulation part, we applied a two-batch setting with sizes (500, 500) and simulated 5 cell groups with group probability (0.2, 0.2, 0.15, 0.2, 0.25). For the second case with “unique cell groups”, we discarded Group 4 and Group 5 cells in Batch 1 and Batch 2, respectively. To generate data sets following model (2), we first fit cFIT with *r* = 5 to obtain a set of 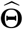. Then we generated our data sets with 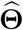 and *σ* = 0.02 in (2), and replaced all negative entries with 0’s. Then the resulting data sets were similar to the original Splatter-generated ones, while possessing a low-dimensional latent structure.

**Table 1:** Simulation settings.

| Simulation setting | Drop-out | #Cells in each batch | Group probability |
| --- | --- | --- | --- |
| base | 0.2 | (500, 900) | (0.4, 0.6) |
| high dropout | 0.33 | (500, 900) | (0.4, 0.6) |
| unbalanced batches | 0.2 | (80, 400) | (0.4, 0.6) |
| unbalanced batches, high dropout | 0.33 | (80, 400) | (0.4, 0.6) |
| multiple cell groups | 0.2 | (500, 500) | (0.3, 0.4, 0.3) |

### 4.5 Analysis of scRNA-seq data

#### Integration of Pancreatic islet cell data sets

The Pancreatic islet cell data sets from five technologies were obtained from R package SeuratData (Lab, 2019), which contains the human pancreatic islet data sets from the following accession numbers: GEO: GSE81076 (CelSeq, 1004 cells), GEO: GSE85241 (CelSeq2, 2285 cells), GEO: GSE86469 (Fluidigm C1, 638 cells), E-MTAB-5061 (SMART-seq2, 2394 cells), and GEO: GSE84133 (inDrops, 8569 cells). The labels provided were treated as the gold standard for visualization. In the pre-processing step, 2000 highly variable genes were selected separately for each data set, followed by a voting procedure to obtain 2000 genes shared by five data sets. Cells from cell types with less than five cells were removed from our analysis. To create the perturbed data sets, we removed all the cells from one cell type in each of the eight data sets. The cell types were chosen as the largest or second-largest cell group in each data set, and selections were alternated so that each cell type were not chosen in more than three out of eight data sets (304 ductal from CelSeq, 844 alpha from CelSeq2, 1008 alpha from SmartSeq2, 258 beta from Fluidigm C1, 868 beta from inDrop1, 659 alpha from inDrop2, 845 acinar from inDrop3, and 104 delta from inDrop4). To identify cell types, unsupervised clustering was performed using the Louvain method on the low dimension representation of integrated data. For cFIT and LIGER result, the estimated factor loading matrices were used, while for Seurat result, PCA was performed to first project data onto a low dimensional space. Then SNN matrix was constructed using the `FindNeighbors` function in Seurat v3 with `k.param` set to 20, followed by identifying clusters using the `FindClusters` command with varying resolution parameters (0.05, 0.08, 0.1). *r* = 15 was used for cFIT that approximated the number of cell types presented in data set.

#### Integration of interneurons and oligodendrocytes from adult mouse

Two data sets of 3212 interneurons and 2524 oligodendrocytes from the mouse frontal cortex (Saunders et al., 2018) were used. The pre-processed scRNA-seq data was retrieved from LIGER repository(Welch et al., 2019). The normalized and scaled data was retrieved and used as input for integration analysis.

#### Integrating data sets of scRNA-seq from developing human brain

Two data sets from the fetal cortex in humans were used for integration. Polioudakis data (Polioudakis et al., 2019) (dbGAP: phs001836) contains 33,976 cells obtained from cortical anlage at mid-gestation (gestation week [GW] 17 or GW18). Samples processed from Drop-seq were obtained from four subjects and two libraries. Using Seurat clustering, the cells were broadly clustered into 16 major cell types by the authors. Nowakowski data (Nowakowski et al., 2017) consists of scRNA-seq of 4,261 cells obtained from multiple brain areas with age varying between 5.85 to 37 post-conception weeks (PCW), and the cells were clustered into 48 subgroups using Seurat and WGCNA by the authors. We pre-processed the two data sets as follows. First, we took the 3790 genes that were identified as enriched in at least one subtype from the authors (Polioudakis et al., 2019) that are also present in Nowakowski data. Then we split the Polioudakis data into five sub-data sets based on the combination of Library and Donor ID and treated each as a separate batch. Then for each data set (including Nowakowski data), UMI counts were normalized by library size and log-transformed (log_2_(*x*/*sum*(*x*) * 10000), and per gene expression was scaled but not centered. The resulting six normalized and scaled data sets were for input into the cFIT integration algorithm (*r* = 40 were used as PCA dimension used in (Polioudakis et al., 2019)). To identify the match cell type of cells in Nowakowski data, we counted the labels of neighboring cells from Polioudakis data (30 nearest neighbors) and labeled the cells by the majority type.

#### Integration of mouse and human MGE cells

To enable the integration across species, human genes were first matched to mouse genes based on HGNC symbols for human genes and MGI symbols for mouse genes (ensemble gene IDs obtained using the `useMart()` function in the biomaRt R package). If multiple matches were identified, averaged expression from multiple matched genes was used.

For mitotic integration analysis, we integrated a data set of mouse MGE cells with human MGE cells. The mouse Single-cell RNA-seq (Drop-seq) was obtained from GSE103983 (Mayer et al., 2018), where 5622 MGE cells from mouse embryos at 13.5 days of gestation (E13.5) were selected. In the original study, three branches were identified through consensus tree construction, which was treated as the gold standard in our analysis to label which trajectory the MGE mouse cells belong. The human MGE cells were taken from Nowakowski data (Nowakowski et al., 2017), including all 733 cells collected from MGE. The authors identified the cells into subtypes of MGE radial glia (MGE-RG), MGE intermediate progenitor cells (MGE-IPC), MGE-div, newborn interneurons (nIN), and other cells that were grouped with other cell types such as striatal interneurons, cortical interneurons derived from CGE and MGE (these cells are labeled ”other” type in our analysis). Human genes were mapped to mouse genes, as described above. 3000 highly expressed genes shared by two data sets were selected. We used r = 10 given limited heterogeneity presented in these cells from the early developmental stage.

For the full mouse and human MGE integration, six data sets were considered. Apart from the mouse Dropseq data from E13.5, four other data sets were added to characterize later developmental stages, including (1) scRNA-seq (10X Genomics Chromium) filtered to 6515 MGE cells (GSE104156). The data set contains only post-mitotic precursors due to the use of FACS sorting to collect GFP-positive cells from MGE dissected from transgenic embryos. (2) scRNA-seq of cortex interneurons (10X) from E18.5 mouse embryos. The data was pre-processed by removing populations of microglia, astrocytes, oligodendrocytes, and smooth muscle cells that probably represent FACS false-positives and are unlikely to give rise to cortical interneurons. The remaining cells were further filtered to the 5568 from MGE eminence. (3) scRNA-seq of cortex interneurons (10X) from mouse at postnatal 10 days (P10), obtained from the same source as E18.5 10X mouse data and subset to MGE. (4) 1350 GABAergic cells (MGE eminence) of P56 adult mouse (Allen Cell Types Database; Tasic et al. (2016)). The pre-processed and filtered version of mouse 10X E18.5 and P10, and Allen P56 data were retrieved from https://www.dropbox.com/s/qe2carqnf9eu4sd/Filtered_Mayer-et-al.Rda.zip?dl=0 (Mayer et al., 2018). For human data, apart from the MGE cells used in the mitotic analysis, we also included 271 cortical interneurons from Nowakovski data (in total 1004 cells from MGE or labeled as IN-CTX-MGE). Human gene expressions were mapped to mouse genes, and 3000 highly variable genes shared among five embryonic data sets were selected. For integration r = 15 was used.

The cFIT R package is available at https://github.com/pengminshi/cFIT.

## Acknowledgments

The authors are grateful to Kevin Lin for helpful comments. This work was supported in part by National Institute of Mental Health (NIMH) grants R01MH123184 and R37MH057881 to K.R. Y.W. was partially supported by the NSF grants CCF-2007911 and DMS-2015447.

